# Variation in glutamate and GABA genes and their association with brain structure and chemistry in autism

**DOI:** 10.1101/2022.05.25.493390

**Authors:** Jilly Naaijen, Martina Arenella, Helge J Zöllner, Nicolaas A Puts, David J Lythgoe, Daniel Brandeis, Sarah Durston, Geert Poelmans, I Hyun Ruisch, Jan K Buitelaar

## Abstract

The excitatory/inhibitory (E/I) imbalance hypothesis posits that an imbalance between glutamatergic and GABAergic neurotransmission contributes to autism symptomatology. Whether this is due to altered GABAergic or glutamatergic functioning, or both, remains largely unknown. We integrated genetic, brain structure and brain chemistry data to investigate the relationship between E/I genetic variation and expression, glutamate concentrations and cortical thickness (CT). Participants (60 autism and 104 neurotypical controls, aged 8-13 years) underwent magnetic resonance imaging and spectroscopy for glutamate quantification in the anterior cingulate cortex (ACC) and left dorsal striatum. Genetic involvement in these regional glutamate concentration levels was investigated using competitive gene-set association and polygenic scores (PGS). Further, glutamate as well as GABA gene-set expression profiles were investigated in relation to CT. Aggregated genetic variation in the glutamate gene-set was associated with ACC but not striatal glutamate concentrations. PGS analysis, however, showed a genome-wide PGS for autism to be predictive of striatal but not ACC glutamate levels. Expression profiles of GABAergic-but not glutamatergic genes were associated with differences in cortical thickness between groups. This study showed differential involvement of aggregated glutamatergic and GABAergic genetic variation in brain structure and chemistry in autism, which suggests regional variability in E/I imbalance.

## 1. Introduction

Autism is characterised by difficulties with social communication and interaction, restricted and repetitive patterns of behavior and/or altered sensory processing (1). The excitatory/inhibitory (E/I) imbalance hypothesis has produced a relatively large body of literature on the neurobiological underpinnings of autism spectrum disorder (autism). According to this almost two decades old hypothesis, dysregulation of the most abundant excitatory and inhibitory neurotransmitters, glutamate and GABA, contributes to autism symptomatology (2). There is increasing evidence supporting the E/I imbalance hypothesis, including animal model work (3,4), the presence of epileptic seizures in people with autism (5), postmortem evidence of altered protein expression (6) as well as the implication of genetic variation associated with glutamate and GABA and changes in their brain concentrations in autism (7–9).

Excitation allows neurons to respond to the environment, while inhibition tunes neuronal selectivity (10). Metabolic evidence for E/I imbalance in autism points to regional variability, but the general direction of the imbalance tends towards cortical disinhibition, especially in childhood (11). Cortical disinhibition, leading to increased excitation, is a common feature within neurodevelopmental conditions such as autism, but also ADHD, schizophrenia and intellectual disability (12). Whether this disinhibition is due to altered GABAergic or altered glutamatergic functioning or both, remains largely unknown.

Although many studies show indirect evidence for an E/I imbalance, there remains inconsistency in the literature. A very recent genetic study investigated the role of genetic variants associated with the GABA_A_ receptor subunit (SNPs within *GABRB3, GABRG3, GABRA5*) and found an association of these with an increased likelihood for autism (13), while the most recent autism GWAS did not find any genome-wide significant associations with glutamate or GABA genes (14). Additionally, a gene-set analysis using the Psychiatric Genomics Consortium (PGC) data did not find an association between autism diagnosis and a glutamate pathway geneset (15). A polygenic score analysis in a Japanese cohort, however, showed associations with autism traits. Further, glutamatergic signalling associated gene-sets were enriched with autism traits and receptive language skills (16).

Animal work is also in support of the E/I imbalance hypothesis: In a recent study using the VPA-induced autism rat model, upregulation of several genes encoding subunits of the ionotropic glutamate receptors was found in the cerebral cortex (17). Altered expression levels of GABAergic genes were found as well, but less pronounced. In humans, gene expression levels have been studied using donors from the Allen Human Brain Atlas (AHBA) (18). Studies up to date have focused on cell-type specific genes according to single RNA sequencing from the mouse hippocampus (19) and investigated whether gene expression across the cortex was related to structural brain variation (e.g. (20–22)). Some genes within these cell-type specific sets refer to cellular E/I function, but only now studies start to focus specifically on glutamate and GABA gene expression throughout the cortex (9).

A different way of investigating glutamate and/or GABA in the brain is by using proton magnetic resonance spectroscopy (^1^H-MRS). Although ^1^H-MRS cannot distinguish neurotransmitter-from overall neurometabolite concentrations, it is the only non-invasive method for in vivo quantification of brain concentrations. Several studies have shown altered glutamate levels in autistic individuals with a focus on regions involved in the cortico-thalamo-striatal cortical (CTSC) loops (e.g. (7,23–26). For GABA, findings are a little more pronounced and point mostly to decreased levels in autism in sensorimotor, frontal and anterior cingulate and cerebellar regions (27–29). For a review of ^1^H-MRS studies into E/I imbalance, see also (26).

In previous studies using a partly overlapping sample, we showed higher glutamate levels in autistic people compared with neurotypically developing controls (NTC) (7) and a reduction in anterior cingulate cortex (ACC) glutamate with age in autistic individuals (8). Here, making use of participants with and without autism from the same sample, we measured multiple aspects of E/I (im)balance by investigating whether brain glutamate concentrations are associated with glutamatergic genetic variation. By using competitive gene-set analysis, we investigated the association between the genetic variability in a carefully selected glutamate gene-set and glutamate concentrations in the ACC and left dorsal striatum. Further, we explored whether the genetic likelihood for autism may be predictive of the concentrations of glutamate in our regions of interest, making use of polygenic scores (PGS). Lastly, we included gene expression data from the open source AHBA to examine whether regional profiles of glutamatergic and GABAergic gene expression across the cortex are associated with structural brain variation in our cohort.

## 2. Methods and materials

### 2.1 Participants

Participants were part of the European TACTICS cohort (www.tactics-project.eu) for whom at least structural MRI data was available and consisted of 60 participants with autism and 104 NTC participants, aged 8-13 years.

For a detailed description of the entire TACTICS sample, see (30). In brief, exclusion criteria for all participants were any contraindications for MRI, an (estimated) IQ < 70 and the presence or history of any neurological disorders. For the NTC participants, no DSM axis I diagnoses were allowed nor in any relatives up to two generations back. Ethical approval for the study was obtained from each study site separately. Parents (and participants > 12 years) gave written informed consent. Diagnosis of autism was informed by preassessment of the Autism Diagnostic Interview Revised (ADI-R; (31)) by a trained researcher. For each step of the analyses, the number of participants differs slightly due to data availability per modality. This is clarified where appropriate and in **Table S1**.

### 2.2 MR acquisition

All MR scans were acquired on 3T scanners. The scanning procedure included a structural T1-weighted MRI scan used for investigation of cortical thickness (CT) as well as for the localization of the spectroscopy voxels for each participant. This scan was matched as closely as possible to the ADNI-GO protocols (32,33). T1-weighted images were further processed in FreeSurfer to extract measures of cortical thickness and for spectral voxel placement and later tissue segmentation for partial volume correction. Proton spectra were acquired using vendor native point resolved spectroscopy (PRESS) sequences, with similar acquisition parameters across sites, using water suppression with chemically selective suppression (CHESS; (34)). Two 8 cm^3^ (2×2×2) voxels were used, one on the midline pregenual ACC and the second on the left dorsal striatum, covering caudate and putamen (**Figure 1**). Following the minimum reporting standard (MRSinMRS) and MRS-Q quality assessment, all acquisition parameters and scans-site specific parameters are shown in detail in **Table S2** (35,36).

**Figure 1.**
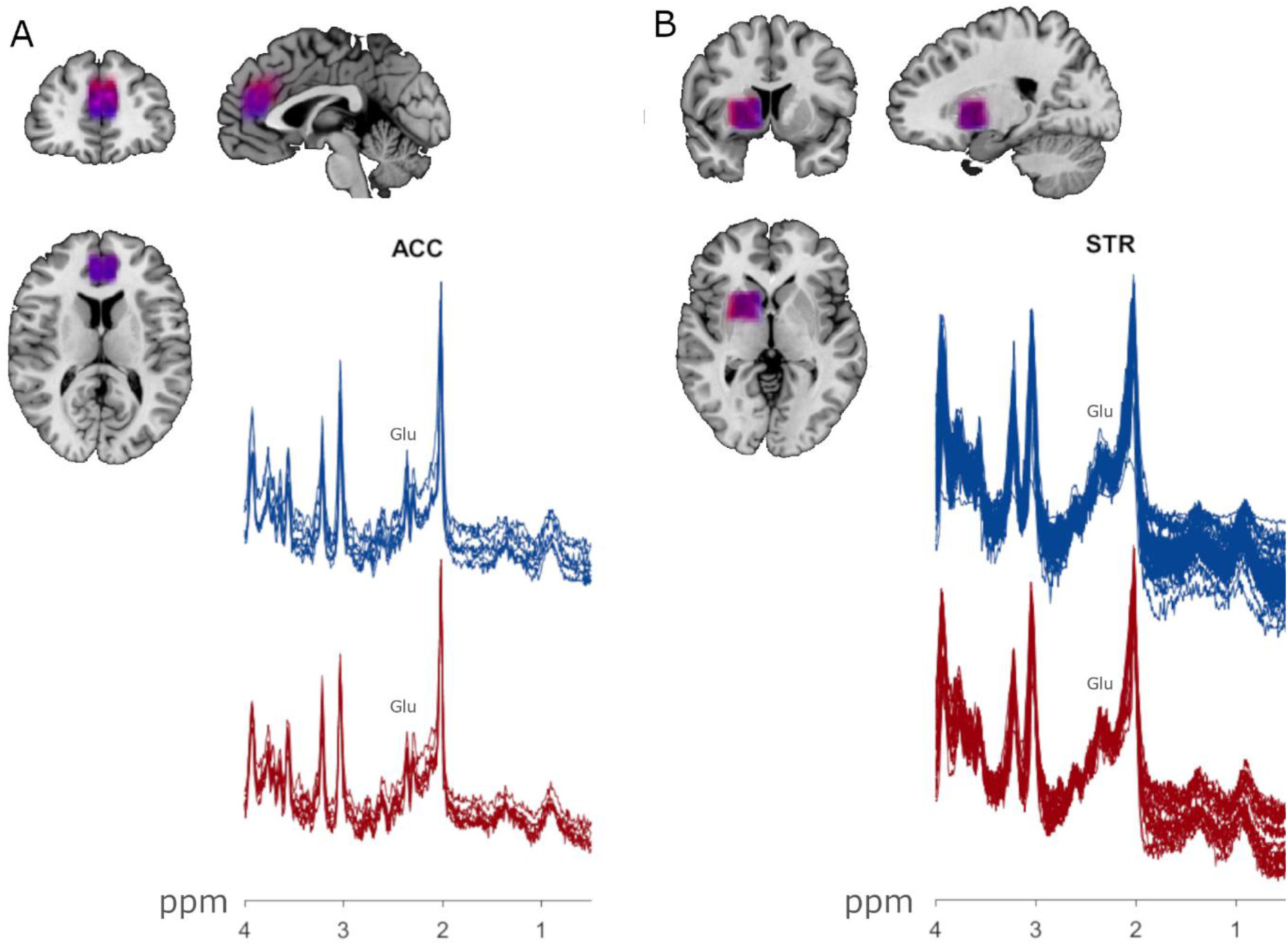
Superposition of average voxels on the MNI152 template and group spectra for the**(A)** midline anterior cingulate cortex and **(B)** left dorsal striatum separately for the groups. Autism = red, NTC = blue. Voxel placement happened consistently across groups as shown by the large overlap and the narrow spatial confinement of the colors. Glu = glutamate peak, ppm = frequency chemical shift in parts per million.

### 2.3 Genetic information

Genome-wide genotyping of the TACTICS participants was performed using the PsychChip_v1-1_15073391 platform in Bonn. Standard GWAS quality control procedures (including filtering based on minor allele frequency (MAF), Hardy-Weinberg equilibrium (*p*-value > 1×10e-6), single nucleotide polymorphism (SNP) call rate (> 95%), subject call rate (> 90%), principal component analysis) and imputation (1000 Genomes reference panel) were performed based on RICOPILI (37). The imputed data underwent additional quality control, in which SNPs with an imputation information score (INFO) lower than 0.8 and MAF lower than 0.05 were excluded. After this step, 5.139.250 SNPs across the autosomal genome were retained (no X-chromosome data available). Genetic data were available for n = 75 NTC and n = 31 autism participants.

Selection of the glutamate and GABA gene-set (74 genes and 128 genes, respectively; **Table 1**) was based on a previous study (38) and updated accordingly to date and using Ingenuity Pathway Analysis (IPA) software (http://www.ingenuity.com). This is a frequently updated genetic database for genetic pathway analyses. The *canonical* pathways described in IPA are based on experimental evidence from the scientific literature and other sources such as gene expression and annotation databases to assign genes to different groups and categories of functionally related genes. This glutamate gene-set was used throughout the association, PGS and gene expression analyses, while the GABA gene-set was used in the gene expression analysis only, due to the unavailability of brain GABA concentrations.

**Table 1.**
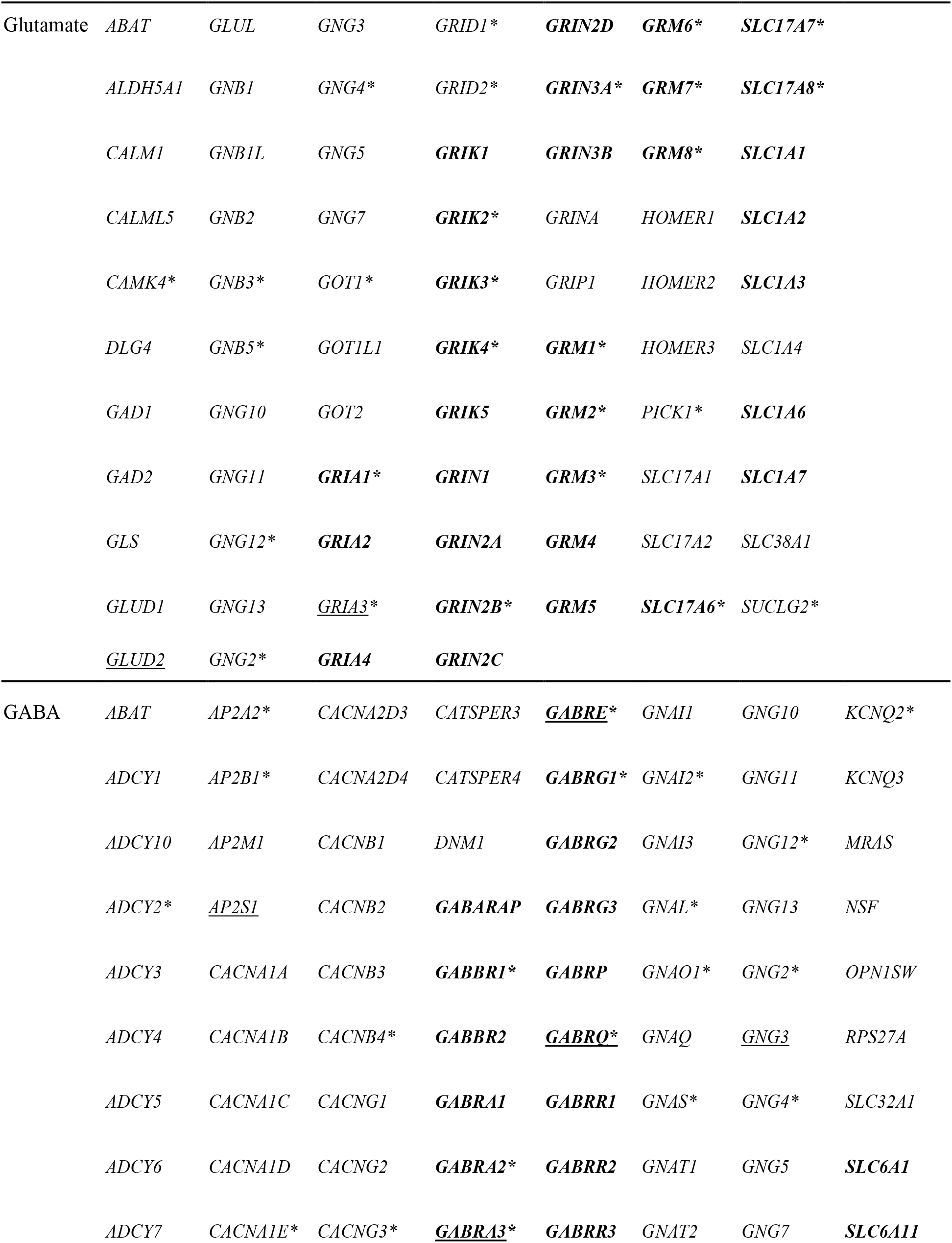
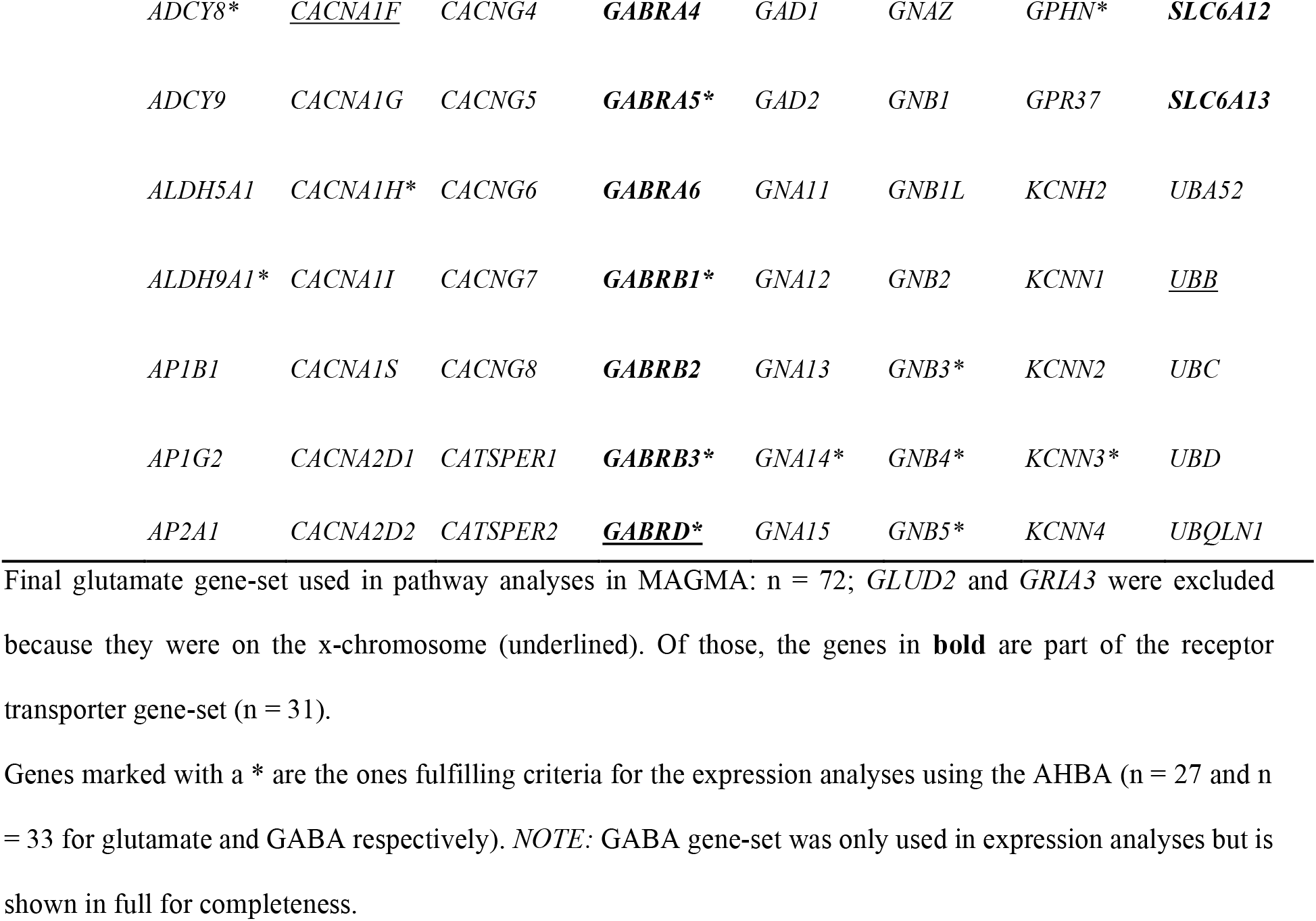
Genes included in the glutamate gene-set (n = 74) and GABA gene-set (n = 128)

### 2.4 Gene expression

Glutamate- and GABA-set gene expression data were obtained from postmortem human brains from the Allen Human Brain Atlas (AHBA; Allen Institute for Brain Science: http://www.brain-map.org (18)), using data from six donors (aged 24-57 years, one female) of the left hemisphere only (right hemisphere data were available for only two donors). Using procedures as described before (20–22) and developed by (39), we mapped the gene expression data from the AHBA to the (left) 34 cortical regions defined by the FreeSurfer in the Desikan-Killiany-atlas (40). To ensure consistency of gene-expression across donors and across the life-span we applied a conservative filtering process. First, a donor-to-median correlation was used to retain only genes with profiles that were consistent across donors (correlation of > 0.46); next, an independent atlas with data across the lifespan (BrainSpan) was used to investigate interregional profile similarity between the AHBA and Brainspan (correlation of > 0.52) (see (39)). This left us with 1.728 genes (out of the 20.737 genes available in the AHBA) to use for the analyses in the current manuscript. For our gene-sets, 27 glutamate and 33 GABA genes were left after this filtering stage, see **Table 1**.

### 2.5 Processing and statistical analyses

#### 2.5.1 Cortical thickness

Cortical thickness (CT) was calculated using FreeSurfer (v 7.1.1) using fully automated and validated procedures, described extensively elsewhere (41,42). For each vertex on the reconstructed cortical sheet, CT was defined as the closest distance between the grey/white matter boundary and the grey matter/cerebrospinal fluid boundary (43). Vertex-wise between group differences in cortical thickness were examined using a linear model with diagnostic group as our variable of interest and age, sex and acquisition site as covariates. For this, we used QDECR (44), a recently developed R package (45,46) for vertex-wise analyses of cortical morphology. Group differences (autism -NTC) in the standard 34 left hemisphere Desian-Killiany atlas regions were used in the gene expression analyses (section 2.5.3).

#### 2.5.2 MRS

Spectroscopy data were processed using OSPREY (47), an open-source Matlab-based software for automated and uniform pre-processing, linear combination modelling, tissue correction and quantification. Vendor-native data were loaded, including metabolite and unsuppressed water reference data. These data were preprocessed into averaged spectra. Eddy-current corrections (48) were performed (where appropriate) using the water reference data. The processed spectra were fitted using OSPREY’s linear combination model with fully localised 2D density-matrix simulated basis functions for each vendor (Siemens, GE, Philips) using the default settings of OSPREY across a frequency range of 0.5 to 4.0 ppm (see **Table S2**). The next steps included coregistration to the participant’s structural image and segmentation of the structural image to generate fractional tissue volumes for each region of interest using SPM12 (https://www.fil.ion.ucl.ac.uk/spm/software/spm12/). Quantification of glutamate was obtained using both water-scaled glutamate concentration estimates corrected for the fraction of CSF as well as tissue-specific relaxation correction according to (49), resulting in molal concentration estimates of glutamate for the ACC and left dorsal striatum. Average spectra for both groups are shown in **Figure 1**. Group differences and associations with symptom measures have been reported in a partly overlapping sample before (7) showing higher levels of ACC glutamate in autism compared to NTC but no differences in striatal glutamate. In this study, we focused on their association with the gene-set only. The ^1^H-MRS methodology allowed us to assess glutamate but not GABA concentrations. From the overlapping sample of genetic and ^1^H-MRS data, a total of 15 spectra had to be excluded due to out of volume artefacts, subject motion or unacceptable linewidth (ACC: 4 (autism = 2 and NTC = 2); left dorsal striatum: 11 (autism = 7 and NTC = 4). See **Table S3** for a comparison between autistic and NTC participants regarding spectral quality and fractional tissue volume after removal of these participants (ACC: 98; striatum: 83).

#### 2.5.3 Genetic analyses

##### Gene-set association

Gene-set association analyses were performed using MAGMA software (version v1.09a; (50)). The 1000 genomes European panel and NCBI 37.3 (retrieved from http://ctglab.nl/software/magma) were used for SNP annotation to genes. All SNPs within the exonic, intronic and untranslated regions of the gene and within 100kb downstream and upstream of the gene were included. After that, we used single gene-based analyses to investigate the degree of association of each gene in the set with each phenotype of interest (so-called gene-based *p*-value, based on multiple linear principal component regression using an *F*-test). Next, we tested the association of the entire gene-set, aggregating the gene-based *p*-values according to their presence in the gene-set by means of a competitive analysis. This tests whether a gene-set is differently associated with the phenotype compared to the remaining genes in the genome, while considering gene size, gene density and linkage disequilibrium (LD) between SNPs.

We investigated whether our glutamate gene-set (74 genes) was associated with glutamate concentrations in the ACC and left dorsal striatum in the sample for whom genetic and spectroscopy data were available after quality control. In case of significant associations, we further explored a smaller gene-set containing only genes encoding receptors and transporters (32 genes) because of their most central role in neurotransmitter metabolism (51), and hence a more likely association with brain concentrations of glutamate.

##### Polygenic scores

In addition to gene-set association, polygenic score (PGS) analyses were performed using PRSice2 software (52). We used the summary statistics of the PGC ASD GWAS (14) as a base dataset and a total of 4.094.303 SNPs could be included for PGS-analyses (SNP matching between the base and our target phenotype was based on rs-numbers). SNPs were clumped based on LD using PRSice default settings (i.e., a bidirectional 250Kb-window and R^2^-threshold of 0.1), resulting in a total of 103.043 LD-clumped SNPs. No prior studies investigating a shared genetic aetiology between autism and levels of brain glutamate have been performed from which we could select an a-priori *p*-value threshold for the PGS. We therefore calculated PGS at multiple *p*-value thresholds using the default PRSice settings to investigate the polygenic signal across a range of SNP-subsets and to avoid under-fitting of the PGS-model (52,53). Multiple testing correction was then applied by calculating an empirical *p*-value for the association of the best-fitting PGS. To this end, the *p*-value of the PGS for the actual phenotype (brain glutamate levels in our target TACTICS sample) was compared with a null-distribution of *p*-values of the PGS regressed on 10.000 randomly permuted phenotypes.

We additionally explored gene-set PGS using the same glutamate gene-set as used in the association testing described above. Here, the autism-PGS is restricted to a limited number of genes present in the set. We computed a competitive *p*-value for the gene-set PGS in relation to the target phenotypes to investigate genetic sharing specifically for the gene-set representing the PGS. The *p*-value threshold for the gene-set PGS was set at 1 since gene-set PGSs containing a small portion of SNPs may be unrepresentative of the whole gene-sets (54).

##### Gene expression

Interregional profiles of glutamate and GABA gene expression, derived from the AHBA, were correlated across the 34 FreeSurfer regions (left hemisphere only) with a profile of group differences (autism minus NTC) in cortical thickness (based on 163 participants; 60 autism/103 NTC), generating a distribution of correlation coefficients for our gene-sets. These distributions were tested for significance using a resampling approach from 10.000 random samples of genes using in-house written scripts in R following the procedures as described in (39).

## 3. Results

The autism and NTC groups did not differ from each other with regard to age, sex and IQ (all *p*-values > 0.05, see **Table 2**). ADI-R scores were only available for the autistic participants.

**Table 2.** Demographic and clinical information from the total number of included participants (n = 164)

|  | **Autism (n = 60)** | **NTC (n = 104)** | **test-statistic** | **p-value** |
| --- | --- | --- | --- | --- |
| Sex m/f | 46/14 | 73/31 | $X^2 = 0.51$ | 0.48 |
| Age, mean (sd) | 10.79 (1.53) | 10.85 (1.32) | $t = 0.27$ | 0.79 |
| IQ, mean (sd) | 107.27 (15.00) | 110.09 (11.47) | $t = 1.33$ | 0.19 |
| ADI-Revised |  |  |  |  |
| Reciprocal interaction | 18.35 (5.31) | - |  |  |
| Communication | 13.53 (3.61) | - |  |  |
| Repetitive behavior | 3.80 (2.57) | - |  |  |

Abbreviations: ADI, Autism Diagnostic Interview (31); m/f, male/female; sd, standard deviation.
*Note:* ADI scores were not available for the NTC group.

### 3.1 Between-Group Differences in Cortical Thickness

Using vertex-wise analysis (QDECR (44)), we did not find any group differences in cortical thickness (all cluster *p*-values > 0.05). Older age was associated with lower cortical thickness in a number of clusters (see **Supplementary material** and **Figure S1**).

### 3.2 Gene-set and brain glutamate

We tested the association of the glutamate gene-set (**Table 1**) with brain glutamate concentrations in the sample that had both genetic and MRS data available (ACC: 98 and striatum: 83). There was a significant association with ACC (*BETA* = 0.167, *SE* = 0.09, *p* = 0.031) but not striatal glutamate (*BETA* = -0.006, *SE =* 0.10, *p* = 0.53) across the entire sample. These results were not driven by one or a few individual genes (all individual gene *p*-values > 0.05). Limiting the ACC results to the receptor/transporter gene-set showed no significant association (*p* = 0.18).

### 3.3 Polygenic Scores

The genome-wide ASD-PGS was linked with striatum glutamate at a *p*-value threshold of 0.004 (best-fitting R^2^ = 0.048, 3100 SNPs, *p* = 0.041), but this did not survive multiple comparison correction using the 10.000 permutations (*p*_*empirical*_*=* 0.35). No association was found with ACC glutamate (best-fitting R^2^ = 0.02, 1905 SNPs, *p* = 0.167, *p*_*empirical*_ = 0.79). We further explored whether the ASD-PGS was predictive of ACC and striatum glutamate concentrations using PGS_set_ including only the glutamate gene-set (74 genes) at a *p*-value threshold of 1. No significant associations were found for the glutamate PGS with ACC (*p* = 0.18) or striatal glutamate (*p* = 0.50).

### 3.4 Gene expression

Interregional variation in the expression of GABAergic genes (33 genes after quality control in the AHBA donor data using (39)) was negatively associated with the interregional profile of group differences in cortical thickness (autism-NTC) and differed significantly from the null-distribution (t = -3.80, *p* = 0.0014, Cohen’s *d* = -1.44). This was not the case for the variation in the expression of glutamatergic genes (27 genes, t = 0.06, *p* = 0.96, Cohen’s *d* = 0.02). Regions with greater expression of GABAergic genes showed a greater difference in CT between autism and NTC. **Figure 2** shows the density plots of the correlation coefficients, variations in group differences in CT and, as an example, the overlap with the variation in expression across the same regions for the most significantly negatively associated gene (Pearsons *r* = -0.28) of the GABA gene-set (*ALDH9A1*, which encodes a protein that is involved in the biosynthesis of GABA). Using total CT in the autism and NTC group separately instead of group differences, we found *positive* associations with the profiles of expression of both the glutamatergic and GABAergic gene-sets (see **Figure S2**), reflecting the not so prominent differences in CT in our cohort as well as a more prominent role for GABAergic genes in cortical thickness in the autism group as compared to the control group.

**Figure 2.**
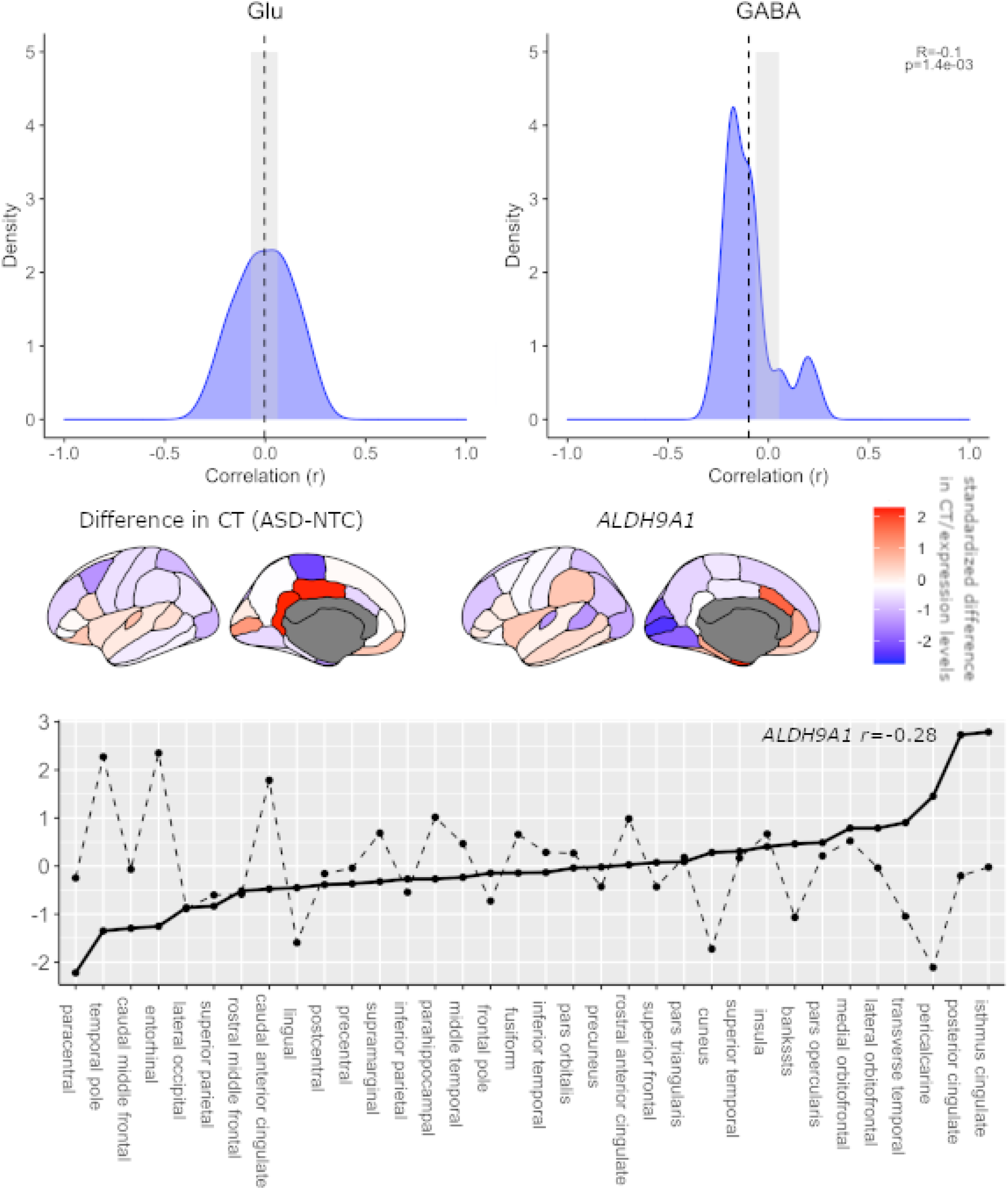
Top-panel. Empirical distribution of the expression-thickness correlation coefficient for the glutamate and GABA gene-set. The x-axis indicates the correlation coefficients between average difference in cortical thickness (autism minus NTC) and expression profile and the y-axis indicates the estimated probability density for the correlation coefficients. The vertical dashed lines indicate the average expression/thickness-difference coefficients across all the genes of the gene-sets; the grey box indicates the 95% confidence interval from the empirical null distribution. The vertical line is outside of the grey box for the GABA-set, indicating the null hypothesis of no association between GABA expression and thickness-difference to be rejected at alpha <0.05. ***Middle panel*** Distributions of the standardised median gene expression levels obtained from the Allen Human Brain Atlas for *ALDH9A1*, which had the strongest negative correlation with the thickness-difference profile (*r*=-0.28). The ranges of the standardised thickness-difference values and expression levels are indicated by the color-scale bar on the right. ***Lower panel***. The plot shows the standardised profiles of the thickness-difference (indicated by a solid line) and the gene expression (indicated by black dotted line). The names of (FreeSurfer) cortical regions are shown on the x-axis in order of thickness level from low to high.

## 4. Discussion

We investigated the E/I imbalance hypothesis of autism using a multimodal approach where we integrated brain glutamate concentrations as measured with MR spectroscopy with genetic variation using both gene-sets and polygenic scores in a cohort of autistic and non-autistic children. To the best of our knowledge, the integration of brain and gene glutamate information has not been performed in autism before. In addition, GABA and glutamate gene expression profiles across the cortex were compared with cortical thickness profiles. Our study shows important and differential roles for glutamatergic and GABAergic genes in relation to autism and brain structure and chemistry. We found genetic variation in the glutamate gene-set to be associated with ACC but not striatal glutamate concentrations. However, polygenic score analyses showed the genetic liability for autism to be predictive of striatal but not ACC glutamate concentrations. Lastly, expression of GABA - but not glutamate -gene-set specific genes was negatively associated with a difference in brain cortical thickness, indicating brain regions with greater expression of these genes to have increased thickness in autistic people compared with controls.

Gene-set association analyses have been widely used in neurodevelopmental disorders, mainly to link aggregated genetic variation to clinical phenotypes (38,55,56). Here we extended this type of analysis by investigating variation in a glutamate gene-set in association with brain glutamate concentrations. We found such association for glutamate concentrations in the ACC but not the left striatum. Although our sample was too small to split this gene-set analysis between the diagnostic groups, we previously showed in the same sample that ACC glutamate was increased in autistic participants ((7), a finding rather robustly shown throughout childhood and adolescence ((57,58). The fact that we did not find glutamate gene-set variation to be associated with left dorsal striatum glutamate concentrations may be due to the fact that the striatum does not contain glutamatergic neurons and receives all its glutamate from cortical afferents (59). We further used polygenic scores to investigate the existence and extent of shared aetiology between autism and brain glutamate concentrations and found a genome-wide polygenic association in relation to striatal but not ACC glutamate. Considering the relatively low *p*-value threshold for the PGS, this indicates that SNPs that associate more strongly with autism may be predictive of increased striatum glutamate concentrations, although this did not survive correction for multiple comparisons. Using a glutamate gene-set specific polygenic score did not reach a similar result, suggesting that the result for the genome-wide PGS does not appear to be exclusively driven by shared glutamatergic genetic variation. Our results indicate differential relationships between glutamatergic genetic variation and ACC and striatal glutamate, respectively, where ACC glutamate concentrations are associated with genetic variation and striatum glutamate concentrations and may share genetics with autism, although especially the latter needs replication in a bigger sample.

Despite many reports of group differences in cortical thickness (60,61), we did not show any significant differences between autistic and control participants in our study. However, we found that regions where GABAergic - but not glutamatergic - genes were increasingly expressed in the normative brain showed larger differences in cortical thickness between our autism and control participants. This was based on gene expression and cortical thickness profiles throughout the cortex. A previous study integrating data from several different conditions showed this negative association as well but with cell-type specific gene expression related to pyramidal cells, astrocytes and microglia (22). Here we specifically focused on genes encoding glutamatergic and GABAergic proteins to pinpoint more precisely to E/I involvement.

In previous studies, links between E/I neurotransmission and autism have been suggested at the genetic level. A study comparing four animal models for autism confirmed an increased E/I ratio due to hyperexcitability and reduced feedforward inhibition (62). A recent systematic review focusing on common genetic variation in autism also confirmed alterations in genes involved in the synthesis of GABA-A receptors as well as both ionotropic and metabotropic glutamate receptors (63). Here we showed that genetic variation was associated with glutamate concentrations in the ACC, a region implicated in autism and related phenotypes (64–66) as well as a region where glutamate concentrations have consistently been reported to be associated with autism before (7,8,23,67). Interestingly, this finding was not confirmed by the PGS analyses. Those showed an association between an ASD PGS and striatal glutamate concentrations. As mentioned above, the striatum in itself contains few glutamatergic neurons, so this association can be speculated to be related to changes in glutamatergic projections from the cortex to the striatum in autism. Although we were not able to measure GABA concentrations, our previous results are in line with increased excitation, which we here integrate with aggregated genetic effects.

Our results should be interpreted in the light of strengths and limitations. One important strength was the integration of E/I measures of brain glutamate concentrations and genetics as well as the addition of expression levels of the gene-sets. Although several studies before have shown changes in glutamate/GABA concentrations throughout the brain and genetic involvement in autism, this was the first integration of ^1^H-MRS and genetic results. However, there are two important limitations to be considered. Firstly, we were not able to measure GABA concentrations in the brain as GABA measurements from PRESS data are not reliable (due to overlap with more highly concentrated metabolites). Thus, our results of the gene-set association and polygenic scores are limited to the excitation part of the E/I imbalance hypothesis. Future studies using more advanced spectroscopic methods can be used to also integrate associations between brain GABA levels and aggregated genetic effects. Secondly, while our sample was fairly large for a ^1^H-MRS study, it was small for a genetic association study and needs replication in a larger sample where differential effects in autism and NTC can be separated for example using ENIGMA MRS (http://enigma.ini.usc.edu/). However, our study can be considered a first proof of concept integrating brain and genetic E/I measures in association with autism, which can be informative for future, larger studies.

In conclusion, we found ACC glutamate concentrations to be associated with aggregated genetic variation in a glutamate gene-set, while polygenic analyses suggested a shared genetic aetiology between autism and striatal glutamate. On the other hand, brain areas where differences in cortical thickness were larger between autism and NTC, showed increased expression of GABA but not glutamatergic genes. These results suggest a complicated, different involvement of glutamatergic and GABAergic genes in autism, and in brain chemistry and structure.

## Acknowledgments

The research leading to these results has received funding from the European Community’s Seventh Framework Program (FP7/2007-2013) TACTICS under grant agreement no. 278948. J Naaijen is supported by a Veni grant from the Netherlands organisation for scientific research (NWO) under grant number VI.Veni.194.032. HJ Zöllner is supported by NIH grants R01EB016089, P41 EB031771, and K99/R00 AG062230. JK Buitelaar has been further supported by the EU-AIMS (European Autism Interventions) and AIMS-2-TRIALS programmes which receive support from Innovative Medicines Initiative Joint Undertaking Grant No. 115300 and 777394, the resources of which are composed of financial contributions from the European Union’s FP7 and Horizon2020 Programmes, and from the European Federation of Pharmaceutical Industries and Associations (EFPIA) companies’ in-kind contributions, and AUTISM SPEAKS, Autistica and SFARI; and by the Horizon2020 supported programme CANDY Grant No. 847818). The funders had no role in the design of the study; in the collection, analyses, or interpretation of data; in the writing of the manuscript, or in the decision to publish the results. Any views expressed are those of the author(s) and not necessarily those of the funders.

The authors would like to thank Saskia de Ruiter, Sophie Akkermans, Nicole Driessen, Vincent Mensen, Bram Gooskens, Muriel Bruchhage, Isabella Wolf, Sarah Hohmann and Regina Boecker-Schlier for their help in data collection, Marcel Zwiers and Paul Gaalman for their technical assistance and all participants and their parents for their participation.

