## Supplementary material for "Variation in glutamate and GABA genes and their association with brain structure and chemistry in autism"

Naaijen et al.

### Data availability

**Table S1.** Available data per modality before and after quality control (QC). The last column shows the numbers of the final analyses.

| Number of participants |  |  |  | Final number included in analyses (autism/NTC) |
| --- | --- | --- | --- | --- |
| Structural MRI: 164 |  |  | → QC → 163 | 60/103 |
| MRS ACC: 151 | Overlapping samples MRS genetic information | ⤴  ⤵ | 102 → QC → 98 | 26/72 |
| Genetic information: 106 |  |  |  | NA |
| MRS Striatum: 128 |  |  | 94 → QC → 83 | 23/60 |

### MRS quality

Table S2 reports the minimal reporting standards for MR spectroscopy (MRSinMRS) studies and Table S3 shows the NAA signal to noise ratio, NAA linewidth in Hz and relative residual amplitude of the linear-combination model (model residual normalised by the noise amplitude) for the ACC and striatum spectra for the autism and NTC group, respectively, showing no difference between the groups regarding spectral quality. Voxel composition differed significantly between groups only for ACC grey matter.

**Table S2.** Minimum reporting standards in Magnetic resonance spectroscopy (MRSinMRS; (35)).

| **Hardware** | | | | |
| --- | --- | --- | --- | --- |
|  | Site 1 | Site 2 | Site 3 | Site 4 |
| Field Strength | 3 T | 3 T | 3 T | 3 T |
| Manufacturer | Siemens | Philips | GE | Siemens |
| Model | Prisma | Achieva | MR750 | Skyra |
| RF coils | 32 channel ^1^H head coil | 32 channel ^1^H head coil | 8 channel ^1^H head coil | 32 channel ^1^H heal coil |
| **Acquisition** | | | | |
| Pulse sequence | | PRESS | | |
| Volume of interest (VOI) locations | | Anterior cingulate cortex (ACC)  Left dorsal striatum | | |
| Nominal VOI size [cm^3^] | | 2 x 2 x 2 cm^3^ (both VOIs) | | |
| Repetition time (T_R_), echo time (T_E_) [ms] | | 3000 / 30 |  |  |
| Total number of excitations or acquisitions per spectrum | | 96 averages  16 water unsuppressed averages | |  |
| Additional sequence parameters | | Spectral width: 5000 Hz  4096 data points | |  |
| Water suppression method | | CHESS |  |  |
| Shimming method | | First- and second order shim | |  |
| Trigger or motion correction method | | None |  |  |
| **Data analysis methods and outputs** | | |  |  |
| Analysis software | | Osprey v 1.1.0 |  |  |
| Processing steps deviating from quoted reference or product | | None |  |  |
| Output measures | | Molal concentrations, water scaled tissue-specific relaxation corrected (49) | | |
| Quantification references and assumptions, fitting model assumptions | | Basis set list: Asc, Asp, Cr, CrCH2, GABA, GPC, GSH, Gln, Glu, Ins, Lac, NAA, NAAG, PCh, PCr, PE, Scyllo, Tau, MM09, MM12, MM14, MM17, MM20, Lip09, Lip13, Lip20  Fitting method: Osprey baseline knot spacing 0.40 ppm | | |
| **Data quality** | |  | | |
| Reported variables (SNR, linewidth) | | Table S3 | | |
| Data exclusion criteria | | Linewidth > 4 x mean linewidth  Lipid contamination  Out of volume signals | | |
| Quality measures of postprocessing model fitting | | Mean relative amplitude residual Tissue fractions  (Table S3) | |  |
| Sample Spectrum | | Figure 1 |  |  |

**Table S3.** MRS quality control for the groups included in the gene-set association analyses.

|  | Autism | NTC | test statistic | *p*-value |
| --- | --- | --- | --- | --- |
| **ACC (n = 98)** | n = 26 | n = 72 |  |  |
| NAA SNR | 222.05 (95.00) | 239.54 (76.07) | t = 0.939 | 0.35 |
| NAA linewidth | 6.39 (3.07) | 6.76 (2.70) | t = 0.587 | 0.559 |
| relresAmp | 9.79 (5.46) | 10.05 (5.58) | t = 0.202 | 0.84 |
| fraction (%)  GM  WM  CSF | 73.73 (3.02)  8.04 (2.61)  18.23 (2.94) | 75.57 (3.06)  7.18 (2.16)  17.25 (2.96) | t = 2.639  t = 1.642  t = 1.455 | 0.01*  0.104  0.149 |
| **Striatum (n = 83)** | n = 23 | n = 60 |  |  |
| NAA SNR | 77.90 (14.41) | 82.15 (14.38) | t = 1.204 | 0.232 |
| NAA linewidth | 15.11 (6.25) | 16.37 (5.23) | t = 0.926 | 0.357 |
| relresAmp | 2.80 (1.02) | 3.03 (1.00) | t = 0.952 | 0.344 |
| fraction (%)  GM  WM  CSF | 62.68 (4.35)  35.99 (4.25)  1.33 (1.58) | 61.02 (6.02)  37.64 (6.03)  1.34 (1.05) | t = 1.221  t = 1.203  t = 0.046 | 0.230  0.230  0.964 |

*Cortical thickness*

Using vertex-wise analysis (QDECR (44)), we found age to be significantly associated with decreasing cortical thickness in a number of clusters, including superior frontal, isthmus cingulate and cuneus regions, depicted in Figure S1 below.

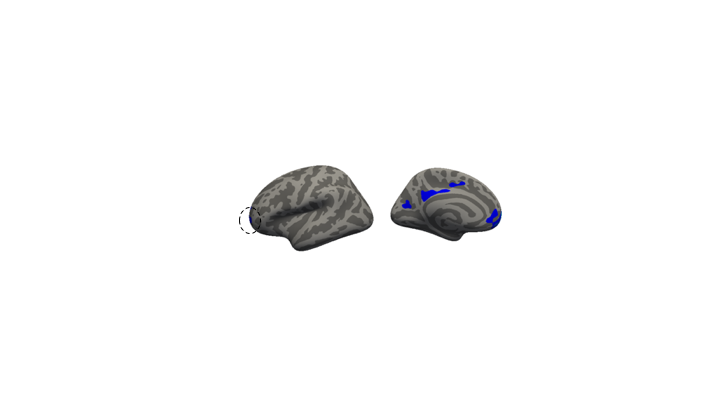

***Figure S1****.* Clusters with significantly lower cortical thickness with older age depicted on the lateral and medial views of the cortex (cluster corrected *p*-values 0.025).

*Gene expression*

Investigating the association between the glutamatergic and GABAergic gene-set expression profiles and cortical thickness (CT) in the autism and NTC group separately, resulted in positive associations for both groups. Figure S2 below shows the density plots for the autism (A) and NTC groups (B).

**A**

**B**
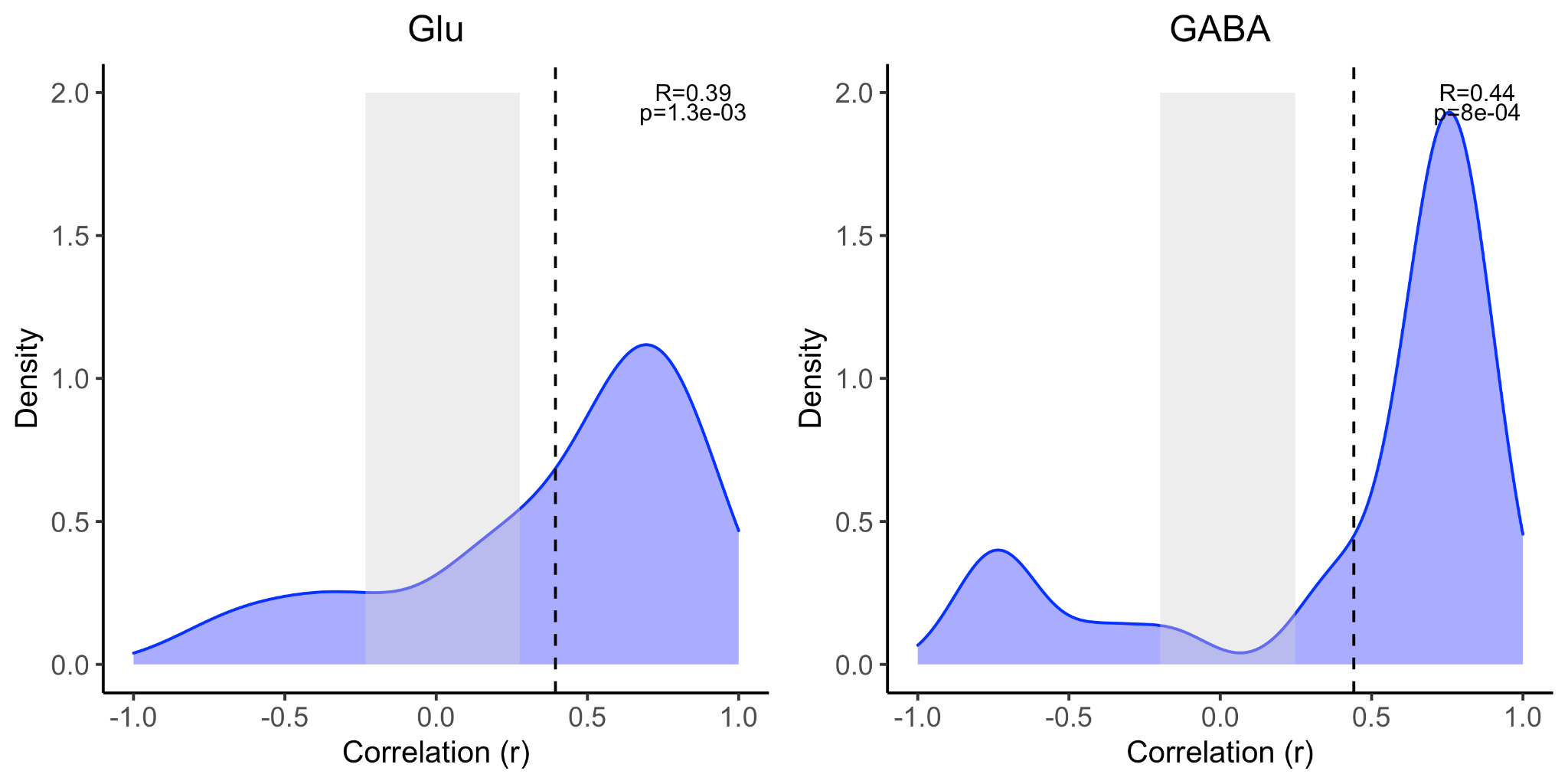

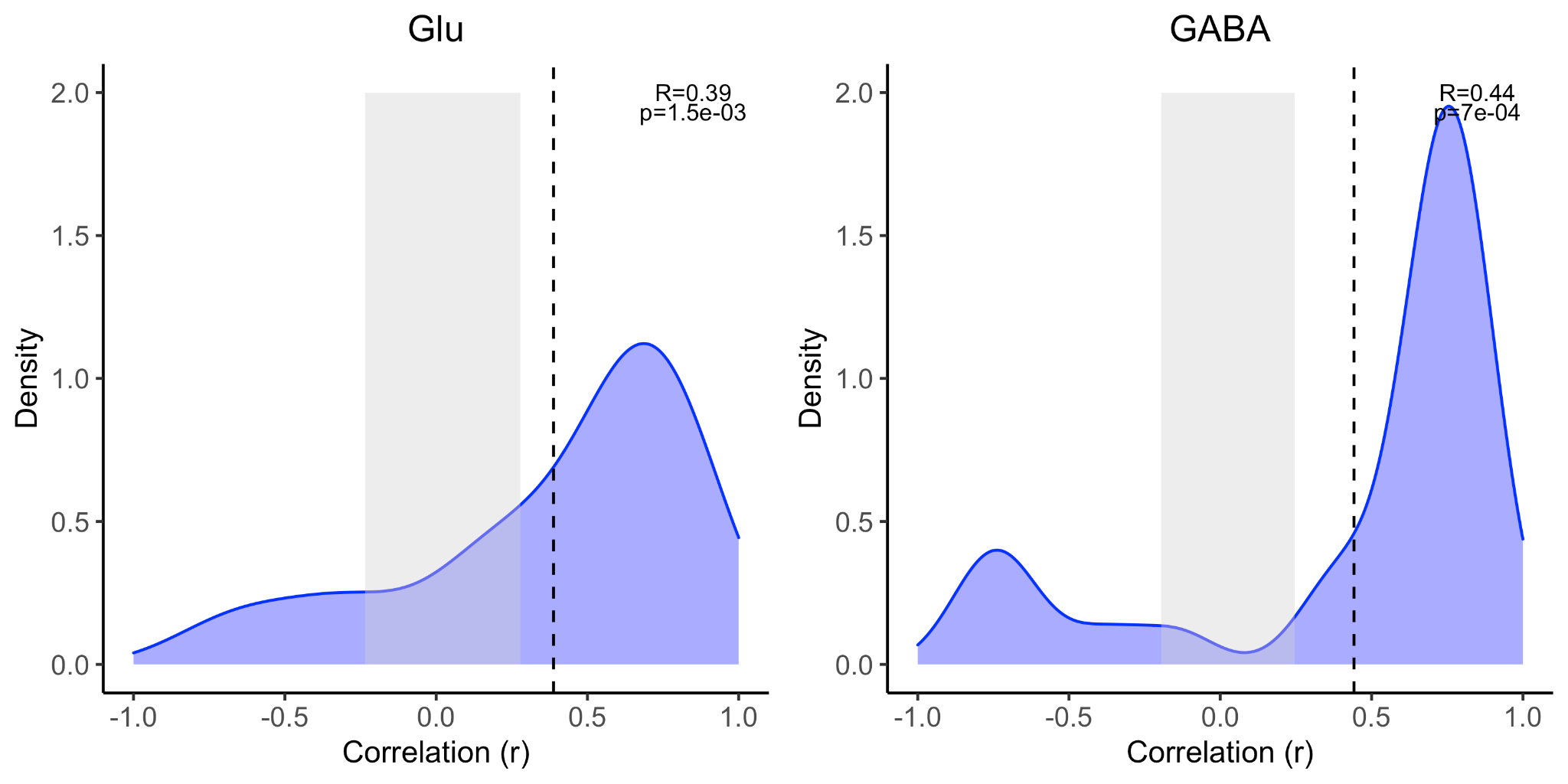

***Figure S2*.** Empirical distribution of the expression-thickness correlation coefficient for the glutamate and GABA gene-set. The x-axis indicates the correlation coefficients between average cortical thickness and expression profile in autism (A) and NTC (B) and the y-axis indicates the estimated probability density for the correlation coefficients. The vertical dashed lines indicate the average expression/thickness coefficients across all the genes of the gene-sets; the grey box indicates the 95% confidence interval from the empirical null distribution. The vertical line is outside of the grey box for both glutamate and GABA in both groups, indicating the null hypothesis of no association between glutamate/GABA expression and thickness to be rejected at alpha <0.05.
